# A new conditional mouse model for Cre-mediated restoration of *Hprt1* expression during brain development

**DOI:** 10.64898/2026.09.15.751558

**Authors:** R.J. Havelaar, A. Fuss, S. Blaess, S.M. Kolk, J.E. Visser

**Author notes:** **Correspondence:** Jasper E. Visser Department of Neurology, Radboud University Medical Center PO Box 9101, 6500HB Nijmegen, The Netherlands *Email:. equal contributions (S.M. Kolk, J.E. Visser).

## Abstract

A central challenge in genetic neurodevelopmental disorders is determining the developmental windows, brain regions, and cell lineages that are affected by gene dysfunction, and whether the resulting pathology remains reversible. The defined genotype–phenotype relationship in Lesch–Nyhan disease (LND), a monogenic disorder caused by pathogenic variants in *HPRT1* and subsequent deficiency of the purine salvage enzyme hypoxanthine-guanine phosphoribosyltransferase (HGprt), provides a well-characterized disease context for such mechanistic investigations.

We present a new conditional *Hprt1* mouse model carrying a loxP-flanked mCherry and transcriptional stop cassette, permitting Cre-driver-dependent analysis of spatially and temporally controlled restoration of endogenous *Hprt1* expression during brain development.

*Hprt1* reactivation was confirmed at the genomic, transcriptional, and biochemical levels. Using the ubiquitous tamoxifen-inducible CAG-CreER driver, a single maternal tamoxifen dose at embryonic day (E)8.5 restored whole-brain *Hprt1* mRNA expression and HGprt enzyme activity in E14.5 embryos to 72.4% and 13.4% of wild-type levels, respectively. Postnatal exposure through lactation during maternal treatment from postnatal day (P)0 to P4 yielded whole-brain HGprt activity equivalent to 28.3% of the WT level at P17. However, ligand-independent CAG-CreER activity and treatment-associated reproductive and developmental complications precluded extended longitudinal rescue studies with this Cre-driver.

Developmentally regulated Wnt1-Cre2 activity, from E8 onwards, provided an alternative approach that permitted analysis through P60. Although the Wnt1 lineage constitutes only a fraction of the developing brain, Cre-dependent reactivation produced detectable HGprt activity in the forebrain, midbrain, and hindbrain. RNA fluorescent *in situ* hybridization (FISH) detected spatially distributed *Hprt1* reactivation in the midbrain and prefrontal cortex at E14.5 and across multiple midbrain, cortical, and subcortical regions at P60.

These findings establish the conditional *Hprt1^tm1.1(LoxP-mCherry-Stop-LoxP)Vsr^*mouse as a model for investigating when and where *Hprt1* restoration could rescue abnormal brain development, while demonstrating that Cre-driver properties determine experimental specificity and longitudinal feasibility. Beyond LND, this conditional reactivation strategy provides an approach for defining developmental windows of reversibility and lineage-specific requirements for gene restoration in monogenic neurodevelopmental disorders.

## Introduction

Lesch–Nyhan disease (LND) is an X-linked neurodevelopmental disorder caused by pathogenic variants in *HPRT1*, the gene encoding the purine salvage enzyme hypoxanthine-guanine phosphoribosyltransferase (HGprt). Near-complete HGprt deficiency causes uric acid overproduction together with severe dystonia, cognitive impairment, and self-injurious behavior.

Clinical observations and experimental models support a developmental rather than degenerative origin of the neurological phenotype. These studies have identified morphological, molecular, and neurochemical abnormalities within dopamine pathways but provide limited information about how and when HGprt deficiency initiates pathology (Bell et al., 2021; Dinasarapu et al., 2022; Göttle et al., 2014; Guibinga et al., 2010; Jinnah et al., 1994; Kang et al., 2011; Lloyd et al., 1981; Witteveen et al., 2022; Wong et al., 1996; Yeh et al., 1998).

Defining these processes and their timing is essential for developing restorative therapies. If neurological dysfunction depends primarily on an ongoing metabolic defect, restoration of HGprt activity – at any time – after disease onset may recover neuronal function. Conversely, if HGprt deficiency during a restricted developmental period irreversibly alters neuronal differentiation or circuit assembly, effective intervention may require restoration within a defined timeframe.

LND is particularly informative for addressing these questions. Its monogenic etiology and defined relationship between residual HGprt activity and clinical severity allow molecular restoration to be linked directly to biochemical and neurological outcomes (Fu et al., 2015). Moreover, the phenotype develops progressively: affected children appear healthy at birth and develop neurological and behavioral manifestations within the first years of life (Jinnah et al., 2006; Mccarthy et al., 2011), while HGprt-deficient mice fail to acquire normal brain dopamine levels during the first postnatal weeks (Jinnah et al., 1994). This extended course provides prenatal and postnatal periods in which to test whether restoring HGprt can redirect an abnormal developmental trajectory.

As constitutive loss-of-function models cannot determine when HGprt is required to prevent pathology or whether established abnormalities remain reversible, an *in vivo* model is required in which endogenous *Hprt1* expression can be restored at defined developmental stages and anatomical locations. Building on our previous developmental studies in *Hprt1*-knockout mice (Witteveen et al., 2022), we introduce a conditional *Hprt1* model carrying a loxP-flanked mCherry reporter and transcriptional stop cassette in intron 1. The cassette disrupts endogenous *Hprt1* expression, whereas Cre-mediated excision restores transcription from the native locus. We validated reactivation at the genomic, transcriptional, and biochemical levels and characterized its temporal and regional performance using CAG-CreER and Wnt1-Cre drivers.

This model enables investigation of when and where *Hprt1* restoration modifies the developmental consequences of HGprt deficiency and provides a proof of principle for conditional gene-reactivation studies that may be applied to other monogenic neurodevelopmental disorders.

## Methods

### Generation of the Hprt1-LSL conditional mouse model

The conditional *Hprt1^tm1.1(LoxP-mCherry-Stop-LoxP)Vsr^* mouse strain (MGI: 6881735) was generated by Cyagen Biosciences Inc. (Santa Clara, CA, USA) using homologous recombination in mouse embryonic stem (ES) cells. A LoxP-mCherry-Stop-LoxP (LSL) sequence were inserted into the first intron of the *Hprt1* gene, downstream of the endogenous *Hprt1* promoter. First, a targeting vector was engineered by generation of the homology arms by PCR using BAC clone RP24-335G16 and RP23-412J16 from the C57BL/6J library as template. A Lox-flanked Neo cassette, self-deleting in germ cells, was used for positive selection, and diphtheria toxin A chain was used for negative selection. Targeted embryonic stem cell clones were injected into C57BL/6 albino embryos, which were then re-implanted into CD-1 pseudo-pregnant females. Chimeric founder animals were identified by coat color. After self-removal of the Neo-cassette, germline transmission was confirmed by breeding with C57BL/6 females and subsequent genotyping of the offspring using primers as listed in Supplementary Table 1. Mice were maintained on a C57BL/6J background. This line is hereafter referred to as Hprt1-LSL.

### Mouse lines, timed mating and genotyping

To evaluate temporally and spatially controlled reactivation of *Hprt1* by Cre-mediated recombination, conditional Hprt1-LSL mice were crossed with CAGGCre-ER^TM^ mice (B6.Cg-Tg(CAG-cre/Esr1*)5Amc/J; The Jackson Laboratory; RRID: IMSR_JAX:004682) or B6 Wnt1-Cre2 mice (B6.Cg-E2f1^Tg(Wnt1-cre)2Sor^/J; The Jackson Laboratory; RRID: IMSR_JAX:022501). CAGGCre-ER^TM^ mice ubiquitously express tamoxifen-inducible Cre recombinase, whereas B6 Wnt1-Cre2 mice express Cre recombinase under the control of the mouse *Wnt1* promoter. The transgenes are hereafter designated as CreER and Wnt1-Cre2, respectively. Sex, *Hprt1* genotype, and Cre-transgene status were determined by PCR analysis of genomic DNA from tail biopsies of embryos or ear punches of postnatal mice. Primer sequences and thermocycling conditions are provided in Supplementary Tables 1 and 2, respectively. PCR products were separated by electrophoresis on 2% agarose gels and visualized with SYBR Safe (Thermo Fisher Scientific).

To ensure unambiguous assignment of developmental stage, females heterozygous for the conditional allele (*Hprt1^LSL/+^*) were mated overnight with Cre-transgenic males (4pm–10am). The morning of vaginal-plug detection was designated embryonic day (E)0.5 and the day of birth was designated postnatal day (P)0. Because *Hprt1* is X-linked and the pathogenic variant is recessive, all studies used male offspring hemizygous for the wild-type or conditional allele. Where possible, wild-type and conditional males were selected from the same litter to optimize age matching and minimize between-group genetic and pre- and postnatal environmental variation.

Males carrying the wild-type *Hprt1* allele (*Hprt1^+^*/Y) were designated WT. Conditional male mutants lacking Cre (*Hprt1^LSL^*/Y) were designated Hprt1-LSL/CreER− or Hprt1-LSL/Wnt1-Cre2− depending on the Cre-harboring line they were crossed with. Conditional mutants carrying Cre, to obtain the recombined *Hprt1^ΔLSL^*/Y allele in Cre-positive cells, were designated Hprt1-LSL/CreER+ and Hprt1-LSL/Wnt1-Cre2+. Where indicated, WT mice were further stratified by Cre status.

Mice were housed in standard cages at 21 ± 1 °C and 60% relative humidity under a 12-h light/12-h dark cycle, with food and water available *ad libitum*. All procedures were approved by the Dutch Central Authority for Scientific Procedures on Animals under project license AVD1030020187065 and conducted in accordance with European, national, and institutional guidelines.

### Tamoxifen treatment

Tamoxifen (Sigma–Aldrich) was prepared in corn oil at 10 mg/mL and administered intraperitoneally, an established route for inducing CreER-mediated recombination *in vivo* (Danielian et al., 1998; Hayashi & McMahon, 2002). For prenatal Cre induction, pregnant dams received a single dose at E8.5 (1 mg/40 g body weight), and embryos were collected at E14.5. For early postnatal induction, dams received tamoxifen once daily from P0 to P4 (4 mg/40 g body weight). This approach has previously been shown to induce recombination in pups through tamoxifen exposure via maternal milk (Silva-Santos et al., 2015; Weber et al., 2009). Offspring was analyzed at P17. Tamoxifen was withheld from animals to assess spontaneous, ligand-independent Cre activity in Hprt1-LSL/CreER+ mice.

### Tissue collection

Following euthanasia of pregnant dams by cervical dislocation, embryos were collected by cesarean section into ice-cold Leibovitz’s L-15 medium (Gibco). Postnatal animals were euthanized by cervical dislocation at the indicated ages. For whole-brain quantitative PCR (qPCR) and enzyme-activity assays, brains were dissected, frozen on dry ice, and stored at −80 °C. For regional enzyme-activity assays in Wnt1-Cre mice, E14.5 brains were grossly divided into forebrain, midbrain, and hindbrain. The posterior border of the cerebral hemispheres was used to define the forebrain–midbrain division, whereas the isthmic constriction, corresponding to the morphological midbrain–hindbrain boundary, defined the caudal limit of the midbrain. The resulting regions were immediately frozen separately on dry ice and stored at −80 °C until analysis.

For histological analysis of Wnt1-Cre2 mice, E14.5 heads and P60 brains were fixed in 4% (w/v) paraformaldehyde (PFA) at 4 °C for 1.5 and 24 h, respectively. Samples were cryoprotected in 30% (w/v) sucrose in phosphate buffered saline (PBS) for 24 h at 4 °C, embedded in M1 embedding medium (Epredia), and stored at −80 °C until sectioning. E14.5 and P60 tissues were cryosectioned at 16 and 20 µm, respectively, using an CM1860 UV cryostat (Leica). Sections were collected on SuperFrost Plus microscope slides (Epredia), dried at 37 °C for 30 min to promote tissue attachment, and stored at −20 °C in sealed containers until immunohistochemical staining.

### RNA isolation, cDNA synthesis and quantitative PCR

Frozen brain tissue was granulated on dry ice using a mortar and pestle, and incubated in 1 mL TRIzol (Invitrogen) for 25 min at 4 °C and 5 min at room temperature (RT). Then, 200 µL Chloroform (ACS grade, 99.8%; Thermo Fisher Scientific) was added, and samples were mixed by inversion and centrifuged at 12,000 × *g* for 15 min at 4 °C. The aqueous phase was recovered, and RNA precipitated with 500 µL isopropanol and collected by centrifugation at 12,000 × *g* for 10 min at 4 °C. Pellets were washed twice with 1 mL ice-cold 75% ethanol, air-dried, and resuspended in nuclease-free Milli-Q water. RNA integrity was assessed by 2% agarose-gel electrophoresis. RNA concentration and purity were measured in duplicate using a NanoDrop ND-1000 spectrophotometer (Isogen Life Science). A total of 500–1,000 ng RNA was treated with DNase I (Thermo Fisher Scientific) according to the manufacturer’s protocol to remove residual genomic DNA. The DNase-treated RNA was subsequently used for cDNA synthesis with the Bioline cDNA Synthesis Kit according to the manufacturer’s protocol.

Quantitative PCR was performed in technical triplicate using a C1000 Touch Thermal Cycler equipped with a CFX96 Real-Time PCR Detection System and operated using CFX MAESTRO software (Bio-Rad). Each 20-µL reaction contained 10 µL of 2 × SensiFAST SYBR master mix (Bioline), 4 µL of 1:15-diluted cDNA, forward and reverse primers at final concentrations of 300 nM each, and nuclease-free water. Primers were designed to span introns where possible. Primer sequences and cycling conditions are provided in Supplementary Tables 1 and 2, respectively. No-reverse-transcriptase controls were included in all experiments. Relative Hprt1 expression was calculated using the ΔΔCt method. ΔCt was calculated as the *Hprt1* Ct minus the mean Ct of *Ppia*, *Actb*, and *Ywhaz*. For each experimental sample, ΔΔCt was calculated by subtracting the mean ΔCt of the corresponding WT group. Expression fold change relative to WT was calculated as 2^−ΔΔCt^. Statistical analyses were performed using individual ΔCt values. For E18.5 analysis of ligand-independent recombination in the CAG-CreER line, three *Hprt1* primer sets targeting distinct transcript regions were used to technically validate the recombination.

### HGprt enzyme activity assay

HGprt activity was measured by conversion of [^14^C]hypoxanthine to [^14^C]Inosine monophosphate (IMP), modified from (Page et al., 1986). Frozen brain tissue was sonicated in assay buffer (50 mM HEPES, 15 mM MgCl₂, pH 7.2) and centrifuged at 14,000 × *g* for 5 min to remove insoluble material. Supernatant (10 µL) was incubated with 40 µL assay buffer containing 25 µM [^14^C]hypoxanthine (Campro Scientific, cat. ARC 0364-50U; 0.1 mCi/mL) and 1.5 mM phosphoribosyl pyrophosphate (Sigma-Aldrich) for exactly 30 min at 37 °C.

Reactions were terminated with 150 µL ice-cold DEAE-Sepharose (Amersham Biosciences), a weak anion-exchange resin that binds negatively charged products, followed by incubation on ice for 10 min. After centrifugation at 14,000 × *g* for 5 min, the resin was washed five times with 500 µL 50% methanol. Resin-bound [^14^C]IMP was eluted with 1 M perchloric acid, followed by centrifugation at 14,000 × *g* for 5 min. Eluate (40 µL) was mixed with 4 mL Opti-fluor scintillation fluid (Perkin Elmer, cat. 6013199) and counted using a Hidex 600SL TDCR counter with a 120-s stabilization time. Counts were corrected against water controls, and negative background-corrected values were set to zero. Protein concentration was determined using the Pierce BCA assay. HGprt activity was expressed as pmol IMP formed/min/µg protein. Relative activity was normalized to the age- and experiment-matched WT mean, while absolute activity values were used for statistical analyses.

### Immunohistochemical staining

Immunofluorescence labeling of mCherry and TH was performed on coronal E14.5 brain cryosections. Sections were rehydrated in PBS for 30 min at RT and blocked for 30 min in blocking buffer (1.67% normal goat serum, 1.67% normal donkey serum, 1.67% normal horse serum, 1% bovine serum albumin, 1% glycine, 0.1% D-lysine, and 0.4% Triton X-100 in PBS). Primary antibody incubation was performed for 16 h at 4 °C in blocking buffer (200 µL/section) containing chicken anti-TH (Abcam, ab76442; RRID: AB_1524535; 1:500) and rat anti-mCherry (Invitrogen, M11217; RRID: AB_2536611; 1:500).Sections were washed three times for 10 min in PBS and incubated for 60 min at RT in the dark with Alexa Fluor 488-conjugated goat anti-chicken IgG (Abcam, ab150169; RRID: AB_2636803; 1:400) and Alexa Fluor 555-conjugated donkey anti-rat IgG (Abcam, ab150154; RRID: AB_2813834; 1:400) in blocking buffer. Sections were washed three times in PBS, counterstained with DAPI for 10 min, washed three additional times, and mounted with Aqua-Poly/Mount (Polysciences). Slides were stored horizontally at 4 °C until imaging.

### RNA fluorescent in situ hybridization (FISH) combined with TH immunofluorescence

To assess spatial patterns of Wnt1-Cre2-mediated *Hprt1* reactivation, multiplex fluorescent RNA *in situ* hybridization for *Hprt1* was combined with immunofluorescent detection of TH on coronal E14.5 (16 µm) and P60 (20 µm) brain sections. Sections were rehydrated in PBS for 5 min at RT, post-fixed in 4% PFA for 5 min, washed in PBS for 5 min and subsequently in PBS containing 0.1% Triton X-100 (PBS-T) for 5 min. After blocking for 1 h in PBS-T containing 10% normal donkey serum, sections were incubated overnight at 4 °C with rabbit anti-TH (Sigma-Aldrich, AB152, RRID: AB_390204; 1:500) in 3% normal donkey serum, washed three times in PBS-T for 5 min, and incubated for 1 h at RT with Alexa Fluor 488-conjugated donkey anti-rabbit IgG (Thermo Fisher Scientific, A21206, RRID: AB_2535792; 1:500). Sections were washed twice with PBS-T for 5 min before proceeding with RNA *in situ* hybridization. RNA was detected using the RNAscope Multiplex Fluorescent Reagent Kit v2 with TSA Vivid Dyes (Advanced Cell Diagnostics/Bio-Techne; 323273), following the frozen-tissue protocol (UM 323100, Rev D, 29/01/2026) with modifications. Probes targeted mouse *Hprt1* (channel C1; 1276311-C1). After immunostaining for TH, sections were dried at 40 °C for 1 h, post-fixed in 4% PFA for 10 min, dehydrated for 3 min each in 50%, 70%, and 100% ethanol, and treated with hydrogen peroxide for 10 min.

E14.5 sections underwent target retrieval at 85 °C for 5 min and Protease Plus treatment at RT for 15 min; P60 sections underwent target retrieval at 100 °C for 3 min and Protease III treatment at 40 °C for 20 min. Probes were hybridized for 2 h at 40 °C, after which sections were washed twice for 2 min in 1 × RNAscope Wash Buffer and stored overnight in 5 × SSC at RT. Signal amplification and channel development were completed according to the manufacturer’s instructions. The *Hprt1* C1 probe was visualized using TSA Vivid 650 (1:3000). Sections were counterstained with DAPI, mounted with Aqua-Poly/Mount (Polysciences), and stored horizontally at 4 °C.

### Imaging and image processing

Immunofluorescence- and FISH-labeled sections were imaged using a Leica DMi8 inverted widefield epifluorescence microscope with a DMC4500 color camera and LAS X software. Tiled z-stacks were acquired using 20×/0.70 NA air, 63×/1.40 NA oil-immersion, or 100×/1.40 NA oil-immersion objectives. Z-stacks were stitched and deconvolved in Huygens Essential software v22–24 (Scientific Volume Imaging), converted to maximum-intensity projections, and exported as TIFF files. Acquisition settings, deconvolution parameters and image-processing steps were kept constant across experimental groups within each experiment. For mCherry quantification, anatomically matched midbrain and prefrontal cortex images were defined using the prenatal mouse brain atlas (Schambra, 2008). Fluorescence intensity was measured in ImageJ v2.16, correcting for background in tissue-free regions.

For quantitative analysis of E14.5 FISH images, anatomically matched 50 × 50-µm fields were selected from the radial and tangential midbrain migration pathways, prefrontal and primary motor cortices, and habenula. *Hprt1* puncta were segmented and counted using a custom Fiji macro with image-specific thresholds and expressed as puncta per µm². Measurements were averaged per animal and region. Field selection and analysis were performed blinded to genotype. P60 FISH data were assessed qualitatively in predefined 100 × 100-µm fields from the midbrain, prefrontal cortex, striatum and habenula. Adult sections were matched rostrocaudally using the Allen Reference Atlas (Dong, 2008), with TH immunoreactivity providing an additional anatomical reference in the midbrain.

### Data analysis and statistics

Statistical analyses were performed in IBM SPSS Statistics v32, and graphs were generated in GraphPad Prism v11. All schematic illustrations, including graphical elements, were created using BioRender.com. The individual animal was the biological unit; technical replicates from each animal and region were averaged before analysis.

To study tamoxifen-induced recombination of the *Hprt1^LSL^* allele after crossbreeding with CAG-CreER mice, E14.5 whole brain *Hprt1* mRNA expression (ΔCt values) and whole-brain HGprt enzyme activities were analyzed by one-way ANOVA, with experimental group as the between-subject factor.

To test for ligand-independent recombination of the *Hprt1^LSL^* allele after crossbreeding with CAG-CreER mice, E18.5 qPCR was analyzed by two-way ANOVA with genotype and Cre status as between-subject factors, and E18.5 and P14 whole-brain HGprt enzyme activities were analyzed by two-way ANOVA with experimental group and age as between-subject factors.

To assess regional recombination of the *Hprt1^LSL^* allele after crossbreeding with Wnt1-Cre2 mice, E14.5 HGprt enzyme activities, mCherry fluorescence and FISH puncta density were analyzed separately by linear mixed-effects models, with genotype as between-subject factor. Brain region was modeled as a repeated measure within subject using a heterogeneous compound-symmetry covariance structure. Models used restricted maximum likelihood estimation and Kenward–Roger degrees of freedom.

HGprt enzyme activities, mCherry fluorescence intensities and FISH *Hprt1* puncta were log2-transformed using Y=log2(Y+0.001) to account for zero values, reduce right skew and stabilize variance. Estimated group differences and 95% confidence intervals were backtransformed and reported as ratios or percentages of geometric means.

Post-hoc comparisons of the estimated marginal means were Šidák-adjusted. Tests were two-sided, with *P* < 0.05 considered statistically significant. Effect estimates are reported with 95% confidence intervals and adjusted *P* values.

End-point allele-specific *Hprt1* PCR and part of FISH results were assessed qualitatively. Data are shown as individual biological observations with mean ± SEM, unless otherwise stated.

## Results

### The *Hprt1* conditional allele couples *Hprt1* disruption to mCherry expression

To permit spatially and temporally controlled reactivation of endogenous *Hprt1*, a genetically modified Hprt1-LSL mouse line was generated carrying the conditional LoF *Hprt1^LSL^* allele. This allele contains a loxP-flanked mCherry reporter and transcriptional stop cassette inserted into intron 1 of *Hprt1* (Fig. 1A). In the absence of Cre recombinase, the cassette disrupts *Hprt1* transcription and permits mCherry expression. Cre-mediated excision of the cassette generates the *Hprt1^ΔLSL^* allele and restores endogenous *Hprt1* transcription.

**Figure 1.**
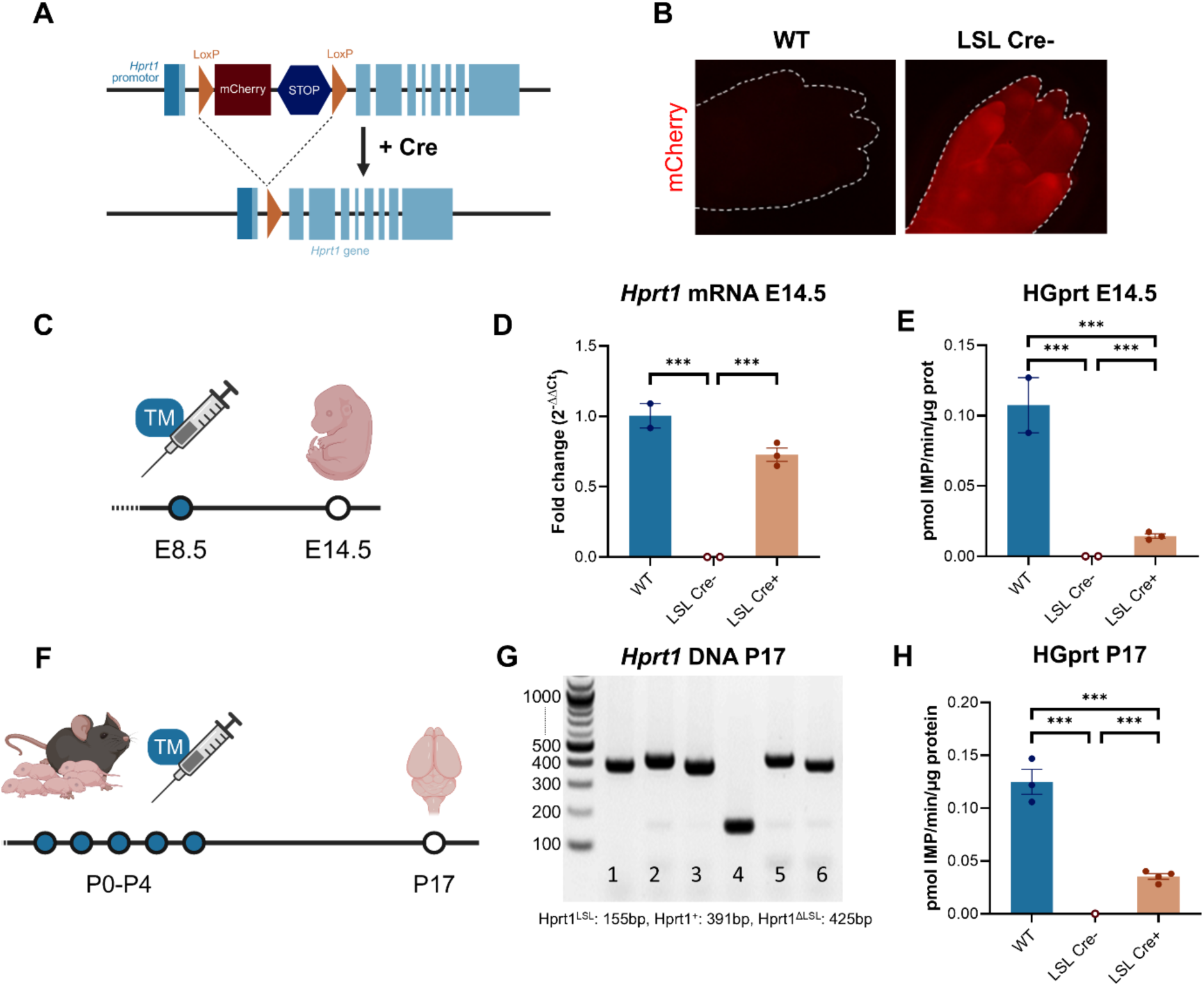
The conditional *Hprt1* allele reports the unrecombined state and permits tamoxifen-induced reactivation. (A) The *Hprt1*^LSL^ allele contains a loxP-flanked mCherry reporter and transcriptional stop cassette in intron 1. Cre-mediated excision generates the *Hprt1*^ΔLSL^ allele, restoring endogenous *Hprt1* transcription. (B) Representative fluorescence images of limbs from Cre-negative WT and Hprt1-LSL embryos at E18.5. (C) Prenatal induction paradigm: heterozygous Hprt1-LSL females (*Hprt1^LSL/+^*) were crossed with CAG-CreER males, dams received a single intraperitoneal tamoxifen injection at E8.5, and male embryos were collected at E14.5. (D) E14.5 whole-brain *Hprt1* expression for WT, Hprt1-LSL/CreER− and Hprt1-LSL/CreER+ embryos, displayed as 2^−ΔΔCt^ relative to WT. (E) Whole-brain HGprt activity (pmol IMP/min/µg/protein) for E14.5 WT, Hprt1-LSL/CreER− and Hprt1-LSL/CreER+ embryos. For D and E, WT, Hprt1-LSL/CreER− and Hprt1-LSL/CreER+ group sizes were *n* = 2, 2 and 3, respectively. (F) Postnatal induction through lactation following daily maternal intraperitoneal tamoxifen injections from P0 to P4, with offspring analyzed at P17. (G) Allele-specific PCR identifying WT (391 bp), *Hprt1*^LSL^ (155 bp) and *Hprt1*^ΔLSL^ (425 bp) alleles. (H) Whole-brain HGprt activity at P17; for WT, Hprt1-LSL/CreER− and Hprt1-LSL/CreER+, group sizes were *n* = 3, 1 and 4, respectively. One-way ANOVA with Šídák-adjusted comparisons was performed on ΔCt values (D) or log₂(Y + 0.001)-transformed activity (E, H). Bars show group mean ±SEM on the original scale; points represent individual animals. *, adjusted *P* < 0.05; **, *P* < 0.01; ***, *P* < 0.001. Abbreviations: WT, wild type; LSL, LoxP-mCherry-Stop-LoxP; Cre, CAG-CreER. Illustrations in A, C and F were created in BioRender. Kolk, S. (2026) https://BioRender.com/p92m2oq https://BioRender.com/5h3ft7i.

Macroscopic mCherry fluorescence was detected under green-light illumination in Hprt1-LSL/CreER− embryos at E18.5 but not in wild-type (WT) littermates (Fig. 1B). These findings support the use of macroscopic mCherry fluorescence to identify embryos carrying the unrecombined *Hprt1^LSL^* allele.

### Tamoxifen-induced Cre activity rescues *Hprt1* transcription and HGprt enzyme activity

We next examined whether removal of the reporter-stop cassette reactivated endogenous *Hprt1* and restored HGprt enzyme function. Hprt1-LSL female mice heterozygous for *Hprt1^LSL^* were crossed with the CAG-CreER line that ubiquitously express tamoxifen-inducible Cre. Pregnant dams received a single tamoxifen dose (1 mg/40 g body weight) at E8.5, and male embryos were collected at E14.5 (Fig. 1C).

Whole-brain qPCR showed negligible *Hprt1* expression in Hprt1-LSL/CreER− embryos. Following prenatal tamoxifen treatment, mRNA expression in Hprt1-LSL/CreER+ embryos reached 72.4% of WT levels (Fig. 1D). Indeed, ΔCt values differed significantly among groups (main effect of genotype, one-way ANOVA, F(2, 4) = 229.30, *P* < 0.001). Compared with Hprt1-LSL/CreER− embryos, mean ΔCt values were 11.4 cycles lower in WT embryos and 10.9 cycles lower in Hprt1-LSL/CreER+ embryos (both Šídák-adjusted P < 0.001). No detectable difference was observed between WT and Hprt1-LSL/CreER+ embryos (mean difference, −0.5 cycles; 95% CI, −2.7 to 1.8; adjusted P = 0.838).

This transcriptional reactivation was accompanied by partial recovery of HGprt activity in Hprt1-LSL/CreER+ whole brains, which reached 13.4% of the WT level (Fig. 1E). Enzyme activity differed among groups (one-way ANOVA, F(2, 4) = 355.5, *P* < 0.001). Post hoc comparisons identified differences between all group pairs (all *P* < 0.001), with HGprt activity in Hprt1-LSL/CreER+ embryos (0,014 pmol IMP formed/min/µg protein) intermediate between that in Hprt1-LSL/CreER− (no detectable IMP formed) and WT (0,107 pmol IMP formed/min/µg protein) embryos.

Thus, a single prenatal tamoxifen treatment administered 6 days before collection at E14.5 restored whole-brain *Hprt1* transcription toward WT levels in Hprt1-LSL/CreER+ male embryos. This transcriptional reactivation was associated with a modest but significant recovery of HGprt activity.

In addition, we tested whether HGprt enzyme activity could also be induced early postnatally by treating nursing dams with tamoxifen from P0 to P4 to transfer tamoxifen through mother milk, a lactational exposure paradigm previously shown to induce CAG-CreER-mediated gene reactivation in neonatal offspring (Silva-Santos et al., 2015; Weber et al., 2009), and analyzing offspring at P17 (Fig. 1F). Allele-specific endpoint PCR detected the recombined allele in tamoxifen-exposed Hprt1-LSL/CreER+ offspring but not in Hprt1-LSL/CreER− controls, confirming stop cassette excision (Fig. 1G).

Applying this treatment regimen, HGprt activity in Hprt1-LSL/CreER+ brains reached 28.3% of WT levels, whereas background-corrected activity in a single available Hprt1-LSL/CreER− brain was nil (Fig. 1H). HGprt enzyme activity differed among groups (one-way ANOVA, *F*(2, 5) = 352.832, *P* < 0.001). Post hoc comparisons identified differences between all group pairs (all *P* < 0.001), with HGprt activity in rescued Hprt1-LSL/CreER+ embryos (0,035 pmol IMP formed/min/µg protein) intermediate between that in Hprt1-LSL/CreER− (no detectable IMP formed) and WT (0,125 pmol IMP formed/min/µg protein) embryos.

In summary, both prenatal and postnatal experiments demonstrate that tamoxifen-induced Cre mediated recombination in Hprt1-LSL mice is able to remove the stop cassette and reactivates the endogenous *Hprt1* locus in developing mice, leading to a partial restoration of HGprt activity as measured in whole brain homogenates.

### CAG-CreER mediates ligand-independent recombination of the *Hprt1^LSL^* allele

Temporal control of gene expression requires efficient induction at the intended time point, while negligible basal expression before induction is equally important. We therefore assessed *Hprt1* expression in Hprt1-LSL/CreER+ animals in the absence of tamoxifen (Fig. 2A, D), to determine whether the system exhibited ligand-independent Cre activity. This distinction is particularly relevant to LND, because residual HGprt activity of only 2–3% is associated with substantial attenuation of the clinical phenotype and absence of the self-injurious behavior characteristic of complete enzyme deficiency (Fu et al., 2014, 2015).

**Figure 2.**
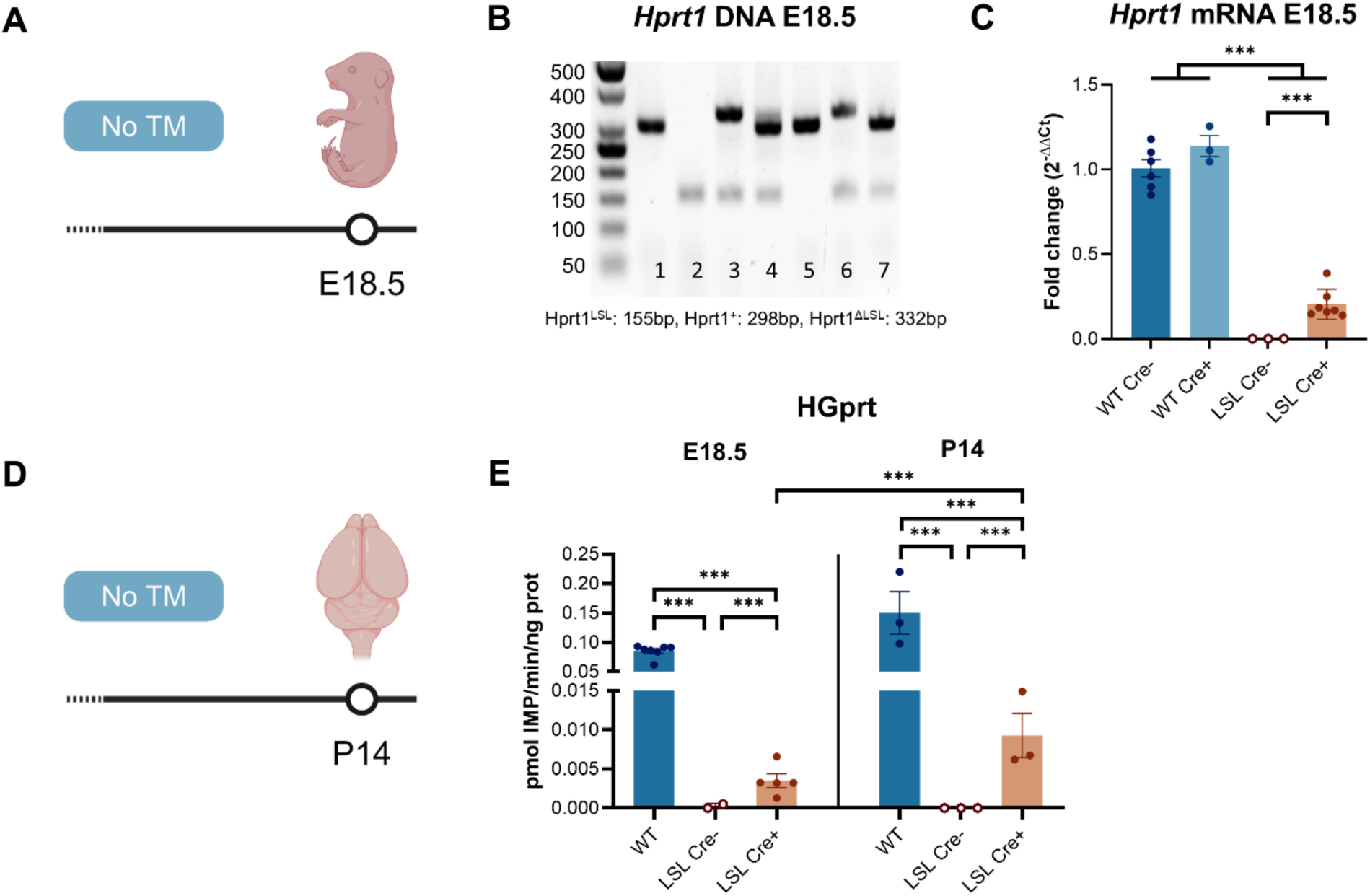
CAG-CreER mediates tamoxifen-independent *Hprt1* reactivation. (A) Experimental design for examining untreated animals at E18.5. (B) Allele-specific PCR identifying WT (298 bp), *Hprt1*^LSL^ (155 bp) and *Hprt1*^ΔLSL^ (332 bp) alleles, including females heterozygous for the conditional allele (*Hprt1^LSL/+^*). (C) E18.5 whole-brain *Hprt1* expression for WT/CreER−, WT/CreER+, Hprt1-LSL/CreER− and Hprt1-LSL/CreER+ embryos, displayed as 2^−ΔΔCt^ relative to WT Cre-. Group sizes were *n* = 6, 3, 3 and 7, respectively. ΔCt values were analyzed by two-way ANOVA with *Hprt1* genotype, Cre status and their interaction, followed by Šídák-adjusted Cre comparisons within genotype. (D) Experimental design for the untreated P14 cohort. (E) Whole-brain HGprt activity (pmol IMP/min/µg protein) at E18.5 and P14, with WT animals pooled across Cre status. WT, LSL Cre− and LSL Cre+ group sizes were *n* = 7, 2 and 5 at E18.5 and *n* = 3 for each group at P14. Log₂(Y + 0.001)-transformed activity was analyzed by two-way ANOVA with experimental group, age and their interaction, followed by Šídák-adjusted group comparisons within age and age comparisons within group. The ages comprised different animals. Bars show group mean ± SEM on the original scale; points represent individual animals. *, adjusted *P* < 0.05; **, *P* < 0.01; ***, *P* < 0.001. Abbreviations: WT, wild type; LSL, LoxP-mCherry-Stop-LoxP; Cre, CAG-CreER. Illustrations in A and D were created in BioRender. Kolk, S. (2026) https://BioRender.com/5h3ft7i.

At E18.5, endpoint PCR detected the recombined *Hprt1^ΔLSL^* allele in all untreated Hprt1-LSL/CreER+ male embryos examined (6/6), but not in Hprt1-LSL/CreER− or WT controls (Fig. 2B). Ligand-independent recombination was also detected in untreated Cre-positive females heterozygous for the conditional allele. Co-detection of the unrecombined *Hprt1^LSL^* and recombined *Hprt1^ΔLSL^*alleles in untreated Hprt1-LSL/CreER+ male embryos was consistent with incomplete or mosaic recombination.

Nevertheless, this genomic recombination was accompanied by substantial transcriptional reactivation. Untreated Hprt1-LSL/CreER+ E18.5 male brains expressed 20.5% of WT *Hprt1* mRNA levels, while Hprt1-LSL/CreER− brains expressed neglectable levels (Fig. 2C). Two-way ANOVA identified main effects of genotype (F(1, 15) = 1626.85, P < 0.001), Cre status (F(1, 15) = 818.26, P < 0.001), and their interaction (F(1, 15) = 767.29, P < 0.001). Indeed, within the conditional mutants, mean ΔCt was 11.41 cycles higher in Cre-negative than in Cre-positive embryos (95% CI, 10.81–12.01; Šídák-adjusted P < 0.001), while Cre status had no detectable effect in WT embryos (mean difference, 0.18 cycles; 95% CI, −0.44 to 0.80; adjusted P = 0.537). Two additional primer sets targeting distinct transcript regions yielded concordant results (Supplementary Fig. 1). Across the three assays, *Hprt1* expression in untreated Hprt1-LSL/CreER+ E18.5 male embryos reached 16–23% of WT levels. These data show substantial ligand-independent transcriptional reactivation in the presence of CAG-CreER at this embryonic age.

### HGprt activity increases with age in untreated Hprt1-LSL/Cre+ mice

To determine whether ligand-independent *Hprt1* reactivation produced detectable HGprt activity, we measured whole-brain enzyme activity in untreated Hprt1-LSL/CreER+ animals at E18.5 and P14. Inclusion of the P14 time point enabled assessment of whether HGprt activity increased with age due to ligand-independent Cre activity (Fig. 2D).

In untreated Hprt1-LSL/CreER+ brains, HGprt activity reached 4.1% of the age-matched WT levels at E18.5 and 6.2% at P14 (Fig. 2E). Two-way ANOVA identified effects of genotype (F(2, 17) = 370.03, P < 0.001), age (F(1, 17) = 7.30, P = 0.015), and their interaction (F(2, 17) = 4.10, P = 0.035). At each age, all pairwise genotype comparisons were significant (all adjusted *P* < 0.001). Post hoc pairwise comparisons for each genotype showed that between E18.5 and P14, HGprt activity further increased by 68% in WT mice (adjusted *P* = 0.028; 95% CI, 7–163%) and by 129% in Hprt1-LSL/CreER+ mice (adjusted *P* = 0.002; 95% CI, 42–269%). Activity in Hprt1-LSL/CreER− mice remained negligible and did not change detectably with age (*P* = 0.48). These findings are consistent with a progressive, age-dependent increase in the functional consequences of ligand-independent Cre activity.

Taken together, the genomic end-point PCR and qPCR data demonstrate tamoxifen-independent reactivation of the conditional *Hprt1^LSL^*allele in Hprt1-LSL/CreER+ embryos and postnatal animals, leading to an HGprt enzyme activity of already 6% in very young mice. Therefore, untreated mice in the current experimental paradigm cannot be regarded as a full HGprt-deficient control.

### Developmental induction limits longitudinal assessment in Hprt1-LSL/CreER mice

Although prenatal induction reactivated the conditional *Hprt1^LSL^* allele, longitudinal follow-up was not feasible. None of the three dams treated with tamoxifen at E8.5 or E12.5 produced viable offspring. The E8.5-treated dam underwent incomplete parturition 23 days after conception. All pups died, and three of the five available for examination had gross abnormalities of the abdomen or posterior body. Both dams treated at E12.5 failed to deliver and were subsequently found dead 25 days after conception. Their uteri contained ten apparently full-term pups without overt gross abnormalities.

Lactation-mediated induction similarly precluded extended follow-up. Three dams received tamoxifen from P0 to P4, indirectly exposing 27 pups that appeared healthy at birth. In the first litter, eight of nine pups died by P29; the remaining pup exhibited severe developmental delay and was euthanized. All pups in the other two litters developed impaired growth, reduced locomotor activity, and diminished stimulus responsiveness, necessitating euthanasia at P21 and P17, respectively. Gross post-mortem examination identified no overt anatomical abnormalities that explained these phenotypes. By contrast, mice administered tamoxifen intraperitoneally at P14 survived to the planned endpoint at P60 without overt complications and were then euthanized for analysis.

Under the conditions tested, neither prenatal nor lactation-mediated induction supported longitudinal rescue studies. Because these observations arose during protocol optimization without prospectively designed vehicle-control groups, the contributions of tamoxifen, vehicle, Cre activity, genotype, maternal effects, and their interactions could not be distinguished. Nevertheless, these outcomes identified feasibility and animal-welfare constraints for extended longitudinal studies involving developmental *Hprt1* induction in the Hprt1-LSL line crossed with CAG-CreER mice. However, the model supported shorter-term *Hprt1* reactivation studies, up to E14.5 and P14 respectively.

### Wnt1-Cre2-mediated recombination restores HGprt activity

Given the limitations of CAG-CreER-mediated temporal control in Hprt1-LSL mice, we investigated developmentally programmed *Hprt1* reactivation as an alternative strategy for longer-term studies. Hprt1-LSL mice were with Wnt1-Cre2 mice to induce recombination in the *Wnt1* lineage and its descendants. In mice, *Wnt1* is expressed between E8 and E14 and is required for dopaminergic midbrain development (Ashly Brown et al., 2011; Stephen Brown & Zervas, 2017; Kim et al., 2021). Although the *Wnt1* lineage constitutes only a relatively small proportion of the brain, we used regional HGprt activity as a functional proof-of-concept readout.

HGprt activity was measured in grossly dissected forebrain, midbrain, and hindbrain from E14.5 WT, Hprt1-LSL/Wnt1-Cre2−, and Hprt1-LSL/Wnt1-Cre2+ embryos (Fig. 3A). In Hprt1-LSL/Wnt1-Cre2+ embryos, activity reached 7.1%, 3.5%, and 2.0% of the corresponding WT levels in the forebrain, midbrain, and hindbrain, respectively. Activity was undetectable in Hprt1-LSL/Wnt1-Cre2− embryos (Fig. 3B).

**Figure 3.**
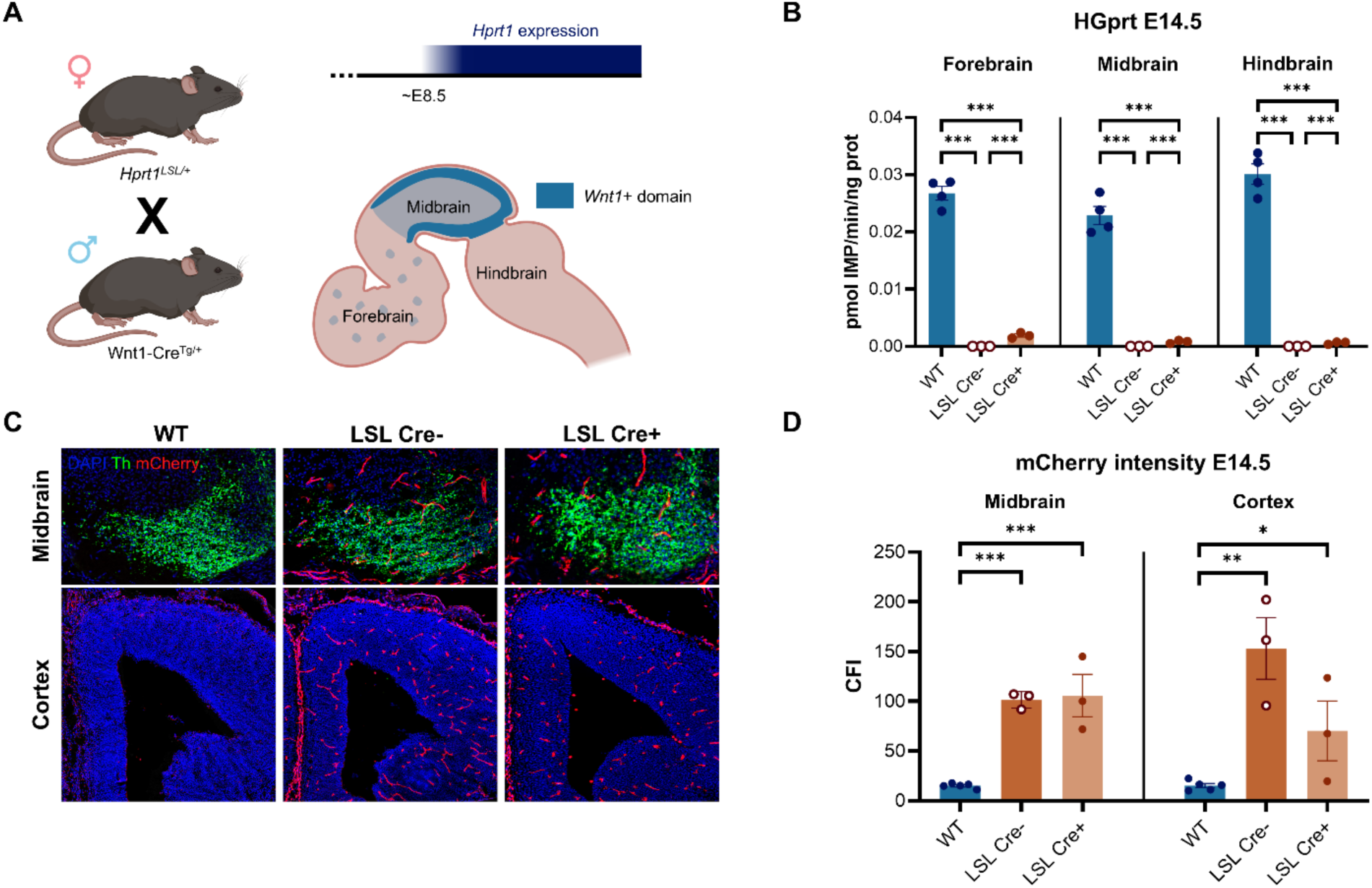
Wnt1-Cre2 restores regional HGprt activity in the embryonic brain. (A) Breeding and developmental reactivation strategy using heterozygous *Hprt1^LSL/+^* females and Wnt1-Cre2^Tg/+^ males. Cassette excision permanently reactivates *Hprt1* in recombined cells and their descendants from E8 onwards. (B) HGprt activity (pmol IMP/min/µg protein) in grossly dissected E14.5 forebrain, midbrain and hindbrain. WT, LSL/Wnt1-Cre2− and LSL/Wnt1-Cre2+ group sizes were *n* = 4, 3 and 3, respectively. (C) Representative DAPI, TH and mCherry immunofluorescence images of the E14.5 midbrain and prefrontal cortex. (D) Corrected regional mCherry fluorescence intensity; group sizes were *n* = 5, 3 and 3, respectively. Panels B and D were analyzed on log₂(Y + 0.001)-transformed values using linear mixed models with group, region and their interaction as fixed effects and region repeated within animal, with heterogeneous compound-symmetry covariance. Pairwise comparisons were Šídák-adjusted. Bars show group mean ± SEM on the original scale; points represent individual embryos. *, adjusted *P* < 0.05; **, *P* < 0.01; ***, *P* < 0.001. Abbreviations: WT, wild type; LSL, LoxP-mCherry-Stop-LoxP; TH, tyrosine hydroxylase; Cre, Wnt1-Cre2. Illustrations in A were created in BioRender. Kolk, S. (2026) https://BioRender.com/2c6k54n.

Linear mixed-effects analysis identified effects of genotype (F(2, 6.67) = 2570.3, P < 0.001), brain region (F(2, 9.54) = 10.65, P = 0.004), and their interaction (F(4, 9.89) = 12.14, P < 0.001). All pairwise genotype comparisons were significant within each region (adjusted P < 0.003), indicating a significant induction of HGprt activity in each region. That these did not reach WT levels was expected, due to the limited fraction of *Wnt1*+ cells in samples. In WT embryos, hindbrain activity was 30% higher than midbrain activity (95% CI, 3-65%; adjusted P = 0.028); the other regional comparisons were not significant. In Hprt1-LSL/Wnt1-Cre2+ embryos, forebrain activity was highest, being 62% higher than midbrain activity (95% CI, 25–110%; adjusted P < 0.001) and 80% higher than hindbrain activity (95% CI, 41-129%; adjusted P < 0.001).

Thus, crossing Hprt1-LSL mice with Wnt1-Cre2 mice reactivates regional *Hprt1* expression and partial recovery of HGprt activity throughout the grossly dissected forebrain, midbrain, and hindbrain.

### Bulk mCherry immunoreactivity to detect *Hprt1^LSL^* to *Hprt1^ΔLSL^* recombination

We next examined whether regional *Hprt1* reactivation was accompanied by loss of the conditional mCherry reporter in E14.5 embryos. Because mCherry immunoreactivity was concentrated predominantly in and around vessel-like profiles, individual labelled cells could not be reliably identified or segmented (Fig. 3C). Reporter abundance was therefore assessed as bulk fluorescence intensity within predefined regions of the prefrontal cortex and midbrain. As expected, no mCherry immunoreactivity was detected in WT embryos, whereas signal remained readily detectable in the prefrontal cortex and midbrain of both Hprt1-LSL/Wnt1-Cre2− and Hprt1-LSL/Wnt1-Cre2+ embryos (Fig. 3C).

Linear mixed-effects analysis identified an effect of genotype (F(2, 8.00) = 43.56, *P* < 0.001), but no effect of region (F(1, 8.00) = 0.41, *P* = 0.542). The genotype-by-region interaction did not reach statistical significance (F(2, 8.00) = 3.45, *P* = 0.083). WT embryos differed from both conditional groups in post hoc comparisons (adjusted *P* = 0.010-0.034). In the cortex, estimated mCherry intensity was 63% lower in Hprt1-LSL/Wnt1-Cre2− than in Hprt1-LSL/Wnt1-Cre2+ embryos, but the difference was not significant (95% CI, −90%–46%; adjusted *P* = 0.175). No numerical difference was observed between the conditional groups in the midbrain (Fig. 3D).

Thus, although Wnt1-Cre2-mediated recombination partially restored HGprt activity in grossly dissected brain regions, bulk mCherry immunoreactivity identified tissue carrying the conditional *Hprt1^LSL^* allele but was insufficiently sensitive to quantify stop-cassette excision at E14.5.

### Wnt1-Cre2 drives spatially distributed *Hprt1* reactivation

To define the spatial pattern of *Hprt1* reactivation in Hprt1-LSL/Wnt1-Cre2+ embryos, we combined FISH detection of *Hprt1* transcripts with immunolabeling for the dopaminergic marker TH. Analyses were performed at E14.5 in the dopaminergic midbrain, given the role of *Wnt1* in midbrain development, and in prefrontal cortical dopaminergic target regions.

We first examined predefined representative 50 × 50 µm fields along the radial and tangential migration pathways of developing dopaminergic neurons at E14.5 (Fig. 4A, B). Qualitatively, WT embryos showed widespread punctate *Hprt1* signals in both regions, in both TH-positive and TH-negative cells. Signal was markedly reduced in Hprt1-LSL/Wnt1-Cre2− embryos but broadly detected in Hprt1-LSL/Wnt1-Cre2+ embryos, without evident restriction to TH-positive cells.

**Figure 4.**
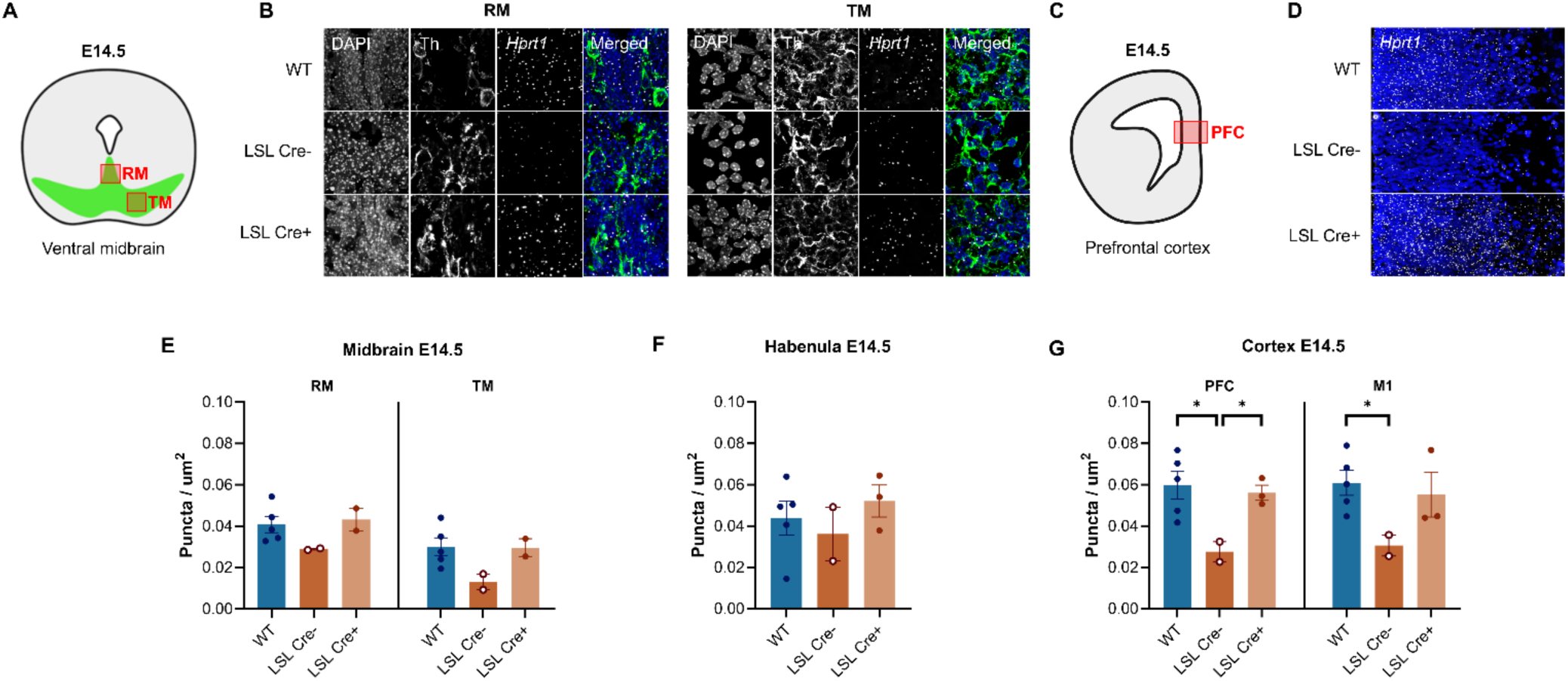
FISH identifies embryonic *Hprt1* reactivation following Wnt1-Cre2-mediated recombination. (A) E14.5 midbrain schematic indicating fields along the radial (RM) and tangential (TM) migration pathways. (B) Representative 50 × 50-µm fields from WT, LSL/Wnt1-Cre2− and LSL/Wnt1-Cre2+ embryos, showing individual DAPI, *Hprt1* and TH channels and merged images. (C) Forebrain schematic locating the prefrontal cortex (PFC). (D) Merged DAPI/*Hprt1* images of 100-µm-wide PFC swatches. (E–G) *Hprt1* puncta density (puncta/µm²) measured in 50 × 50-µm fields from midbrain RM/TM (E), habenula (F), and PFC and primary motor cortex (M1; G). In the group order listed above, sample sizes were *n* = 5, 2 and 2 for RM/TM and *n* = 5, 2 and 3 for habenula, PFC and M1. All five regions were analyzed together on log₂(Y + 0.001)-transformed values using a linear mixed model with group, region and their interaction as fixed effects and region repeated within animal, with heterogeneous compound-symmetry covariance. Group comparisons were Šídák-adjusted. Bars show group mean ± SEM on the original scale; points represent individual embryos. *, adjusted *P* < 0.05; **, *P* < 0.01; ***, *P* < 0.001. Abbreviations: WT, wild type; LSL, LoxP-mCherry-Stop-LoxP; TH, tyrosine hydroxylase; Cre, Wnt1-Cre2. Illustrations in A and C were created in BioRender. Kolk, S. (2026) https://BioRender.com/llca7cl.

A similar pattern was observed in the developing prefrontal cortex. *Hprt1* signal was widespread in WT embryos, sparse in Hprt1-LSL/Wnt1-Cre2− embryos and broadly restored in Hprt1-LSL/Wnt1-Cre2+ embryos (Fig. 4C, D). Thus, Wnt1-Cre2-mediated reactivation was not confined to the sampled midbrain pathways.

To quantify the spatial distribution of *Hprt1* transcripts, puncta density was measured in the radial and tangential midbrain migration pathways, habenula, prefrontal cortex, and primary motor cortex (Fig. 4E–G). The cortical regions were selected as downstream targets of midbrain dopaminergic projections, whereas the habenula was included as a major upstream modulator of dopaminergic activity. Across these regions, mean puncta density was approximately 1.7-fold higher in Hprt1-LSL/Wnt1-Cre2+ than in Hprt1-LSL/Wnt1-Cre2− embryos (0.047 versus 0.027 puncta/µm²). The mean density in Hprt1-LSL/Wnt1-Cre2+ embryos was numerically similar to that in WT embryos (0.047 puncta/µm²).

Linear mixed-effects analysis identified effects of experimental group (F(2, 7.19) = 7.93, *P* = 0.015) and region (F(4, 12.89) = 9.82, *P* < 0.001), without group-by-region interaction (F(8, 13.95) = 1.00, *P* = 0.477). Across regions, estimated puncta density was 41% lower in Hprt1-LSL/Wnt1-Cre2− than in WT embryos (95% CI, −63–-6%; adjusted *P* = 0.028) and 43% lower than in Hprt1-LSL/Wnt1-Cre2+ embryos (95% CI, −66–-4%; adjusted *P* = 0.035). No detectable difference was observed between Hprt1-LSL/Wnt1-Cre2+ and WT embryos (95% CI, −32%–55%; adjusted *P* = 0.997).

Residual puncta in Hprt1-LSL/Wnt1-Cre2− embryos may reflect low-level residual transcription or assay background. Because these signals were not further characterized, their origin cannot be determined definitively. Together with the regional HGprt activity measurements, these findings support spatially distributed transcriptional reactivation of the conditional *Hprt1^LSL^* allele and partial biochemical recovery in Hprt1-LSL/Wnt1-Cre2+ embryos.

### Wnt1-Cre-mediated *Hprt1* reactivation persists into adulthood

Having established Wnt1-Cre-dependent *Hprt1* reactivation at E14.5, we next examined whether expression remained detectable at P60. Exploratory qualitative FISH analysis covered predefined regions of the midbrain, striatum, prefrontal cortex and habenula (Fig. 5A–H). In WT midbrain tissue, abundant *Hprt1* puncta were observed in the ventral tegmental area and substantia nigra pars compacta. Signal varied between individual cells but was present in both TH-positive and TH-negative populations, without obvious qualitative enrichment in either. Relative to WT, Hprt1-LSL/Wnt1-Cre2− tissue showed sparse signal, whereas Hprt1-LSL/Wnt1-Cre2+ tissue exhibited widespread puncta in both midbrain regions (Fig. 5B).

**Figure 5.**
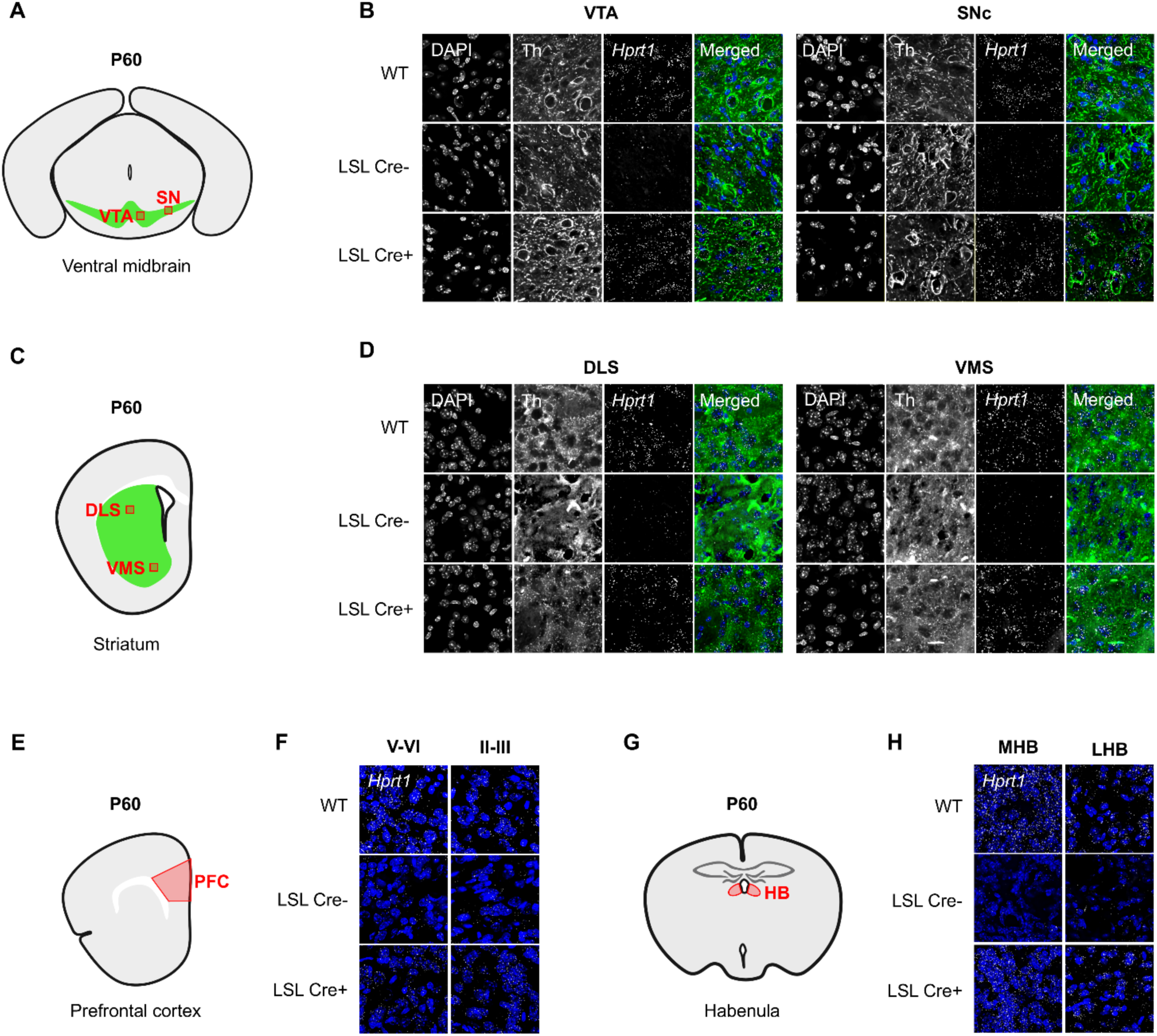
Wnt1-Cre2-mediated *Hprt1* reactivation persists into adulthood. Exploratory qualitative FISH images from P60 WT, LSL/Wnt1-Cre2− and LSL/Wnt1-Cre2+ brains (*n* = 1 animal per group). (A) Midbrain schematic indicating the ventral tegmental area (VTA) and substantia nigra (SN). (B) Images of the VTA and substantia nigra pars compacta (SNc). (C, D) Striatal schematic and images of the dorsolateral (DLS) and ventromedial striatum (VMS). (E, F) Prefrontal cortex (PFC) schematic and images from deep (layers V–VI) and superficial (layers II–III) cortical layers. (G, H) Habenular schematic and images of the medial (MHB) and lateral habenula (LHB). In B and D, columns show DAPI, *Hprt1*, TH and the three-channel merge, respectively. Panels F and H show DAPI/*Hprt1* merges. All image fields are 100 × 100 µm. Abbreviations: WT, wild type; LSL, LoxP-mCherry-Stop-LoxP, TH, tyrosine hydroxylase; Cre, Wnt1-Cre2. Illustrations in A, C, E and G were created in BioRender. Kolk, S. (2026) https://BioRender.com/llca7cl.

A similar pattern extended to the dorsolateral and ventromedial striatum (Fig. 5D), the deep and superficial layers of the prefrontal cortex (Fig. 5F), and the medial and lateral habenula (Fig. 5H). Across these regions, WT tissue showed widely distributed *Hprt1* puncta, whereas signal remained sparse in Hprt1-LSL/Wnt1-Cre2− tissue and was again widespread in Hprt1-LSL/Wnt1-Cre2+ tissue.

Although these analyses were qualitative, they establish transcript presence and anatomical distribution of *Hprt1* expression. Moreover, they show that Wnt1-Cre2-mediated *Hprt1* expression remained detectable at P60, and was widespread present throughout the brain.

## Discussion

In this study we validated, at molecular and biochemical levels, a novel mouse model that allows Cre-mediated restoration of *Hprt1* expression. In the *Hprt1^tm1.1(LoxP-mCherry-Stop-LoxP)Vsr^* mouse, a LoxP-mCherry-Stop-LoxP (LSL) sequence was placed upstream of the *Hprt1* gene, downstream of the endogenous *Hprt1* promoter. In these Hprt1-LSL mice, this *Hprt1^LSL^* allele prevents *Hprt1* transcription and permits mCherry expression instead. Cre-mediated excision of the cassette, to generate the *Hprt1^ΔLSL^*allele, restored endogenous *Hprt1* transcription as well as HGprt enzyme activity, in both embryonic and early postnatal mouse brains. In addition to these proof-of-concept studies demonstrating the feasibility of HGprt restoration in this conditional genetic mouse model, we identified Cre-line specific constraints associated with the CAG-CreER mouse line that must be considered when applying this model for temporally controlled rescue during development.

### *Hprt1* expression and HGprt enzyme activity is rescued by Cre-mediated recombination in the Hprt1-LSL mouse

Reactivation of *Hprt1* expression in Hprt1-LSL mice could be demonstrated at successive biological levels, both pre- and postnatally: mCherry marks the unrecombined *Hprt1^LSL^*allele without *Hprt1* expression, allele-specific endpoint PCR confirms stop-cassette excision after Cre-mediated conversion into the *Hprt1^ΔLSL^*allele, qPCR demonstrated restoration of endogenous *Hprt1* mRNA transcription and the enzyme assay demonstrated biochemically active HGprt in rescued animals.

Using the ubiquitous tamoxifen-inducible CAG-CreER driver, a single maternal tamoxifen dose at E8.5 restored whole-brain *Hprt1* mRNA expression and HGprt activity in E14.5 embryos to 72.4% and 13.4% of wild-type levels, respectively. Postnatal exposure through lactation during maternal treatment from P0 to P4 yielded whole-brain HGprt activity equivalent to 28.3% of the WT level at P17. Thus, a substantial transcriptional recovery translated into a modest enzymatic restoration.

Several mechanisms may explain the disparity between restored *Hprt1* transcript levels and HGprt enzyme activity. For example, Cre-mediated excision retains a single loxP sequence within the 5ʹ untranslated region (UTR). Because 5ʹ-UTR sequence and secondary structure regulate mRNA stability and translation initiation, the residual loxP site could reduce translational efficiency by introducing a stable hairpin that impedes ribosomal scanning (Cautereels et al., 2024; Leppek et al., 2018; Wiles et al., 2014).

Alternatively, limited enzymatic recovery could reflect reduced HGprt catalytic competence rather than reduced protein abundance. Cre-mediated recombination can generate unexpected splice products (Yang et al., 2009), while recombinant HGprt variants can exhibit impaired catalytic properties or stability (Fu & Jinnah, 2012). Distinguishing reduced protein abundance from impaired enzyme function requires characterization of full-length *Hprt1* transcripts and HGprt protein quantification alongside enzyme activity.

More generally, transcript abundance and enzyme activity are not directly comparable because qPCR and biochemical assays differ in normalization and dynamic range. Post-transcriptional regulation and protein turnover further weaken correspondence between mRNA and protein abundance, especially in heterogeneous brain tissue (Bauernfeind & Babbitt, 2017; Schwanhäusser et al., 2011). Developmental profiling of HGprt expression and activity will be required to distinguish these possibilities.

Despite incomplete biochemical recovery, the HGprt activity achieved may be sufficient to modify LND-related neuropathology. In patients, residual enzyme activity is strongly associated with clinical severity: approximately 3% activity has been associated with absence of self-injury, whereas approximately 10% may prevent most neurological manifestations (Fu et al., 2014, 2015; Hersh et al., 1986). The enzyme activity restored in the Hprt1-LSL model could therefore attenuate the dopaminergic abnormalities observed in HGprt-deficient mice, although this requires direct phenotypic evaluation.

Bulk mCherry fluorescence distinguished Hprt1-LSL from WT tissue but did not reliably discriminate *Hprt1^LSL^* form *Hprt1^ΔLSL^* alleles. Persistent mCherry protein following cessation of reporter transcription may have obscured the loss of expression – particularly as tissue was analyzed within days of induction. Moreover, reporter fluorescence does not necessarily provide a quantitative measure of target-locus recombination or function (Fernández-Chacón et al., 2019; He et al., 2019). Accordingly, under the conditions tested here, mCherry fluorescence should be interpreted as a qualitative indicator of conditional-allele carriage rather than a quantitative measure of *Hprt1* reactivation

### Limitations of the CAG-CreER model

Use of CAG-CreER to reactivate *Hprt1* in our studies revealed two limitations: substantial ligand-independent recombination and adverse reproductive and developmental outcomes associated with the induction protocol.

In untreated Hprt1-LSL/CreER+ embryos, ligand-independent recombination restored whole-brain *Hprt1* expression to 16–23% of WT levels by E18.5. HGprt activity subsequently reached 6% of WT levels at P14. Reactivation was confined to Cre-positive animals, implicating the driver–target combination rather than intrinsic instability of the conditional allele. Ligand-independent recombination has been reported for the CAGGCre-ER^TM^ line and other CreER drivers, with its frequency and extent determined by driver activity and target-locus susceptibility (Álvarez-Aznar et al., 2020; Hayashi & McMahon, 2002). Because cassette excision is irreversible, sporadic recombination permanently reactivates *Hprt1* in affected cells and their descendants, compromising temporal control.

This background activity is particularly relevant to LND studies because low residual HGprt activity may substantially attenuate disease severity and the underlying neuropathology, as explained earlier. However, as activity measured in patients’ fibroblasts cannot be directly compared with that measured in mouse brain homogenates (Fu et al., 2015), the phenotypic consequences of ligand-independent *Hprt1* reactivation *in vivo* requires direct evaluation.

Tamoxifen exposure imposed an additional constraint. Reproductive failure, fetal abnormalities, and postnatal developmental delay occurred during protocol optimization. Although the absence of controlled comparator groups precludes attribution exclusively to tamoxifen, these findings are consistent with reported effects of prenatal tamoxifen exposure, including embryotoxicity, structural abnormalities, intrauterine hemorrhage, impaired parturition, and pup mortality (Lizen et al., 2015; Savery et al., 2020; Sun et al., 2021; Ved et al., 2019). These side effects particularly confound the interpretation of developmental outcomes.

### Wnt1-Cre2 drives persistent *Hprt1* reactivation across brain development

Developmentally regulated Wnt1-Cre2 activity, from E8 onwards, provided an alternative approach that permitted analysis through P60. Although the *Wnt1* lineage constitutes only a fraction of the developing brain, Cre-dependent reactivation produced detectable HGprt activity in the forebrain, midbrain, and hindbrain. FISH detected spatially distributed *Hprt1* reactivation in the midbrain and prefrontal cortex at E14.5 and across multiple midbrain, cortical, and subcortical regions at P60, indicating that embryonically initiated reactivation persisted into adulthood.

The distribution of reactivation was broader than anticipated from the established role of *Wnt1* in midbrain development. Germline recombination caused by ectopic Cre expression in male germ cells has been reported (Dinsmore et al., 2022), but is unlikely to explain the present findings because male offspring inherited the X-linked *Hprt1^LSL^*allele maternally and the Cre transgene paternally.

Instead, the broad distribution may reflect embryonic *Wnt1* expression beyond the midbrain (Parr et al., 1993; Wilkinson et al., 1987). Neuroepithelial recombination has been reported with the original Wnt1-Cre driver (Jacques-Fricke et al., 2012), while lineage tracing with the same Wnt1-Cre2 driver used here revealed low-level scattered labeling in the forebrain at E9.5 (Keuls & Parchem, 2021).

These findings therefore demonstrate the feasibility of Wnt1-Cre2-associated reactivation of *Hprt1*, but do not establish that recombination was restricted to the canonical domains of endogenous *Wnt1* expression.

### Strengths and limitations

A principal strength of the presented Hprt1-LSL model is the restoration of *Hprt1* expression from its endogenous locus. Reactivation can be evaluated sequentially through cassette excision, transcript production, and HGprt activity. Compatibility with different Cre drivers and persistence of reactivation into adulthood support its use across developmental stages. Moreover, allele-specific endpoint PCR, qPCR, FISH, reporter imaging, and enzyme assays provide complementary measures of recombination and function.

Several factors limit the resolution of the present study. Small group sizes limited statistical power, particularly in the tamoxifen-induction experiments. Whole-brain and regional homogenates may obscure effects restricted to discrete cell populations. Finally, the CAG-CreER and Wnt1-Cre2 experiments show that temporal and anatomical specificity depends on the interaction between the Cre driver and target allele, rather than on the nominal driver profile alone. Each driver–target combination therefore requires empirical validation.

## Conclusions and future applications

This study establishes the Hprt1-LSL mouse model as a platform for Cre-dependent reactivation of endogenous *Hprt1 in vivo*. Genomic, transcriptional, and biochemical analyses confirmed that cassette excision restores *Hprt1* expression and HGprt activity, although the efficiency and specificity of reactivation depended on the Cre driver.

The model can be used to define when HGprt activity is required for normal dopaminergic development and whether established abnormalities remain reversible. Reactivation before the first detectable abnormality would test prevention, induction during its emergence would assess disease modification, and later restoration would test reversal. The proliferation and migration abnormalities identified in HGprt-deficient dopaminergic progenitors at E14.5 provide a biologically defined reference point for these experiments (Witteveen et al., 2022). Future studies will require tightly regulated Cre drivers and parallel assessment of recombination, *Hprt1* expression, HGprt activity, and relevant neurobiological phenotypes. This approach should help define the cellular requirements and therapeutic windows for HGprt restoration – or any other therapeutic intervention – in LND.

Beyond LND, this conditional reactivation strategy provides an approach for defining developmental windows of reversibility and lineage-specific requirements for gene restoration in monogenic neurodevelopmental disorders.

## Author Contributions

Havelaar RJ: Methodology, Validation, Formal analysis, Investigation, Writing — original draft, Visualization. Fuss A: Methodology, Investigation, Writing — review & editing. Blaess S: Methodology, Writing — review & editing, Supervision. Kolk SM: Writing — review & editing, Supervision, Funding acquisition. Visser JE: Conceptualization, Methodology, Writing — review & editing, Supervision, Project administration, Funding acquisition.

## Acknowledgements

This work was supported by grants from the LND Famiglie Italiane Odv, the National Institute for Neurological Diseases and Stroke (NINDS) at the National Institutes of Health (NIH, R01 NS119758) and the University of Pennsylvania Orphan Disease Center in partnership with the Love Never Sinks Organization to JEV, as well the Dutch Research Council (NWO) for the ProMiSe NWA project (NWA.1160.18.320) to SMK.

The authors thank the personnel at the Animal Research Facility (CDL, Radboud University Medical Center, Nijmegen, the Netherlands) for expert animal care and technical assistance. In addition, Jeroen van de Heuvel (Department of Pharmacy, Pharmacology and Toxicology, Radboud University Medical Center, Nijmegen, the Netherlands) is acknowledged for his assistance in the isotope-labelled HGprt enzyme assays.

## Supplementary informa>on

**Supplementary Table 1.**
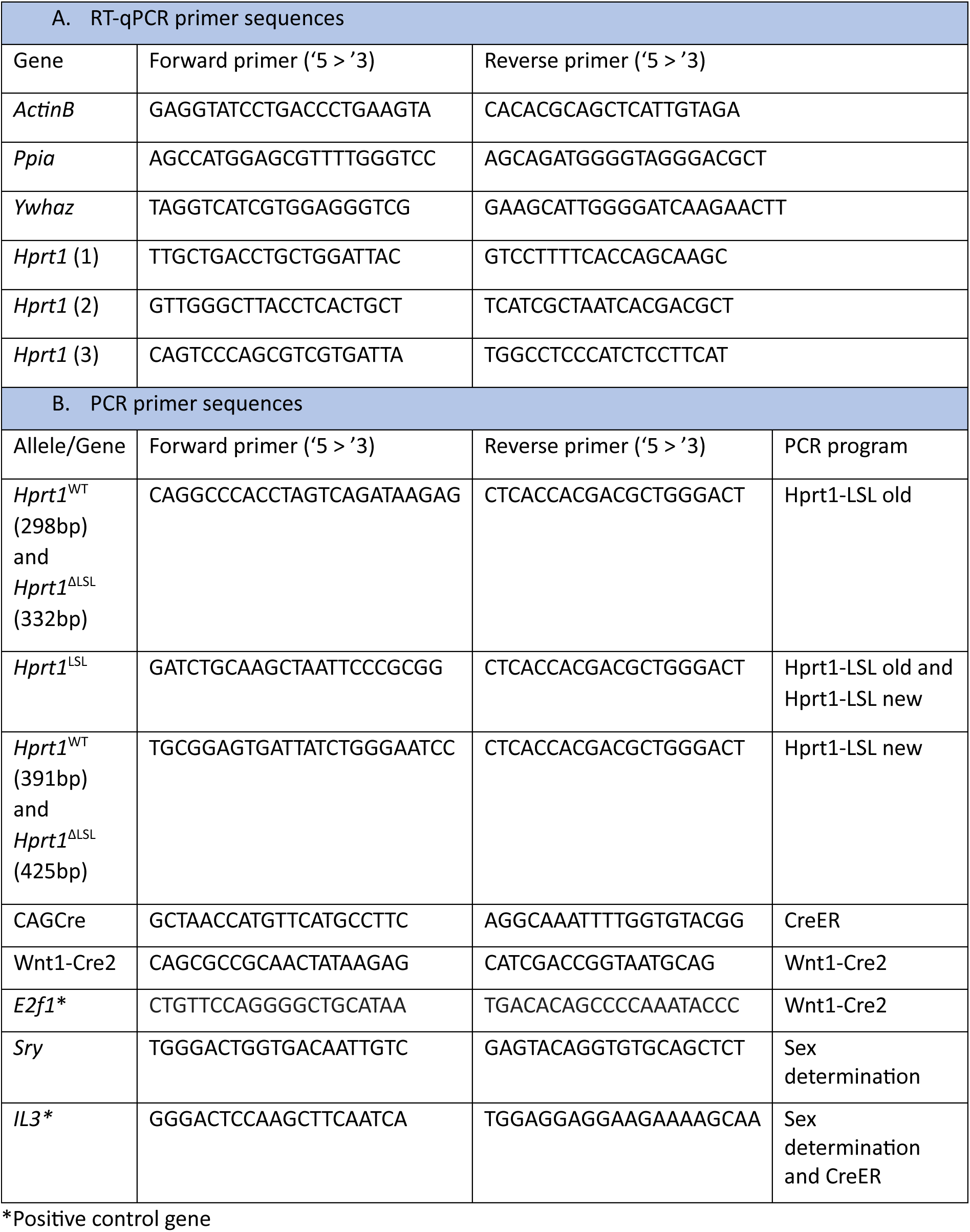
PCR primer sequences.

**Supplementary Table 2.** PCR programs.

a. PCR program Hprt1-LSL old
| temp | Time (min:sec) | cycles |
| --- | --- | --- |
| 95,0 °C | 3:00 | 1x |
| 95,0 °C | 0:30 | 35X |
| 59,6 °C | 0:30 |  |
| 72,0 °C | 0:45 |  |
| 72,0 °C | 5:00 | 1X |
| 4,0 °C | ∞ | 1x |

b. PCR program Hprt1-LSL new
| temp | Time (min:sec) | cycles |
| --- | --- | --- |
| 95,0 °C | 3:00 | 1x |
| 95,0 °C | 0:30 | 35X |
| 63,0 °C | 0:30 |  |
| 72,0 °C | 0:45 |  |
| 72,0 °C | 5:00 | 1X |
| 4,0 °C | ∞ | 1x |

c. PCR program Wnt1-Cre2
| temp | Time (min:sec) | cycles |
| --- | --- | --- |
| 95,0 °C | 3:00 | 1x |
| 95,0 °C | 0:30 | 10X |
| 65-0.5°C* | 0:30 |  |
| 68,0 °C | 0:45 |  |
| 94,0 °C | 0:30 | 30X |
| 60,0 °C | 0:30 |  |
| 72,0 °C | 0:45 |  |
| 72,0 °C | 5:00 | 1X |
| 4,0 °C | ∞ | 1x |

\* Touchdown protocol starting at 65 with 0.5 °C decrease every cycle down to 60.5 °C.

d. PCR program CreER
| temp | Time (min:sec) | cycles |
| --- | --- | --- |
| 95,0 °C | 1:00 | 1x |
| 95,0 °C | 0:30 | 35X |
| 58,0 °C | 1:00 |  |
| 68,0 °C | 0:45 |  |
| 68,0 °C | 5:00 | 1X |
| 4,0 °C | ∞ | 1x |

e. Sex determination
| Temp | Time (min:sec) | Cycles |
| --- | --- | --- |
| 95,0 °C | 03:00 | 1 |
| 95,0 °C | 00:30 | 40 |
| 58,0 °C | 00:30 |  |
| 72,0 °C | 00:45 |  |
| 72,0 °C | 10:00 | 1 |
| 4,0 °C | ∞ | 1 |

f. RT-qPCR
| Temp | Time (min:sec) | Cycles |
| --- | --- | --- |
| 95,0 °C | 02:00 | 1 |
| 95,0 °C | 00:05 | 40 |
| 60,0 °C | 00:20 |  |
| 65,0-95,0 °C* | 00:05 |  |
| 4,0 °C | ∞ | 1 |

\* Melt-curve analysis starting at 65 °C with 0.5 °C increase every 5 seconds up to 95 °C.

**Supplementary Figure 1.**
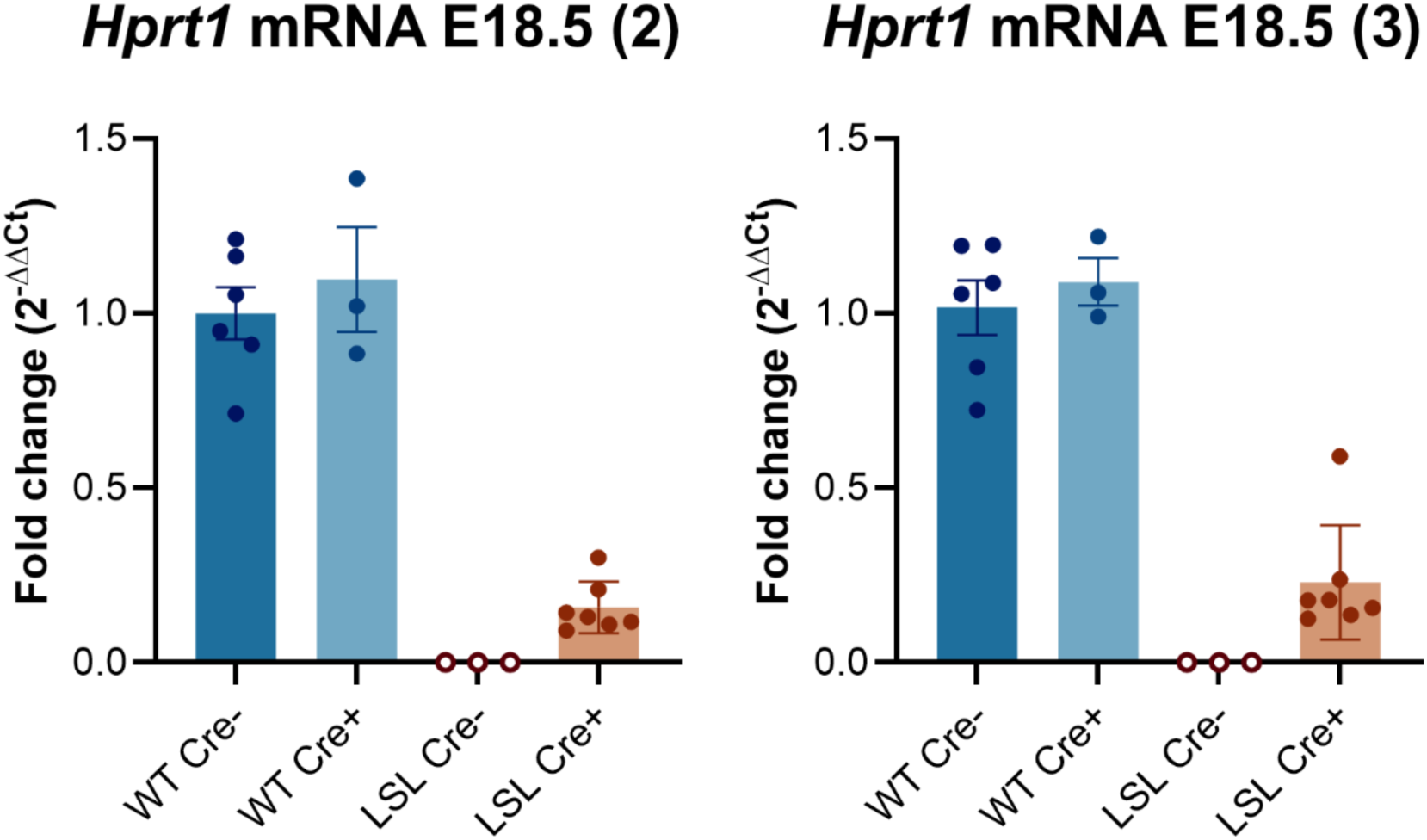
Additional qPCR assays support tamoxifen-independent *Hprt1* reactivation. Whole-brain *Hprt1* expression in untreated E18.5 animals measured using two additional primer sets targeting distinct transcript regions. Expression is displayed as 2^−ΔΔCt^ relative to WT Cre−. Group sizes were WT/CreER−, *n* = 6; WT/CreER+, *n* = 3; LSL/CreER−, *n* = 3; and LSL/CreER+, *n* = 7. The same animals were analyzed with both validation assays and the primary assay in Figure 2C; these assays provide technical validation rather than independent biological replication. Bars show group mean ± SEM; points represent individual animals. WT, wild type; LSL, LoxP-mCherry-Stop-LoxP; Cre, CAG-CreER.

